# Automatic adaptation of model neurons and connections to build hybrid circuits with living networks

**DOI:** 10.1101/419622

**Authors:** Manuel Reyes-Sanchez, Rodrigo Amaducci, Irene Elices, Francisco B. Rodríguez, Pablo Varona

## Abstract

Hybrid circuits built by creating mono- or bi-directional interactions among living cells and model neurons and synapses are an effective way to study neuron, synaptic and neural network dynamics. However, hybrid circuit technology has been largely underused in the context of neuroscience studies mainly because of the inherent difficulty in implementing and tuning this type of interactions. In this paper, we present a set of algorithms for the automatic adaptation of model neurons and connections in the creation of hybrid circuits with living neural networks. The algorithms perform model time and amplitude scaling, drift compensation, goal-driven synaptic and model tuning/calibration and also automatic parameter mapping. These algorithms have been implemented in RTHybrid, an open-source library that works with hard real-time constraints. We provide validation examples by building hybrid circuits in a central pattern generator. The results of the validation experiments show that the proposed dynamic adaptation facilitates building hybrid circuits and closed-loop communication among living and artificial model neurons and connections. Furthermore contributes to characterize system dynamics, achieve control, automate experimental protocols and extend the lifespan of the preparations.

## 1 Introduction

Hybrid circuits are networks built by connecting model neurons and synapses to living cells. Pioneering works in building such interactions go back almost three decades ago (Yarom, 1991) with many successfull implementation since then, e.g. see (Szücs et al, 2000; Pinto et al, 2000; Varona et al, 2001; Le Masson et al, 2002; Nowotny et al, 2003; Olypher et al, 2006; Arsiero et al, 2007; Grashow et al, 2010; Brochini et al, 2011; Wang et al, 2012; Hooper et al, 2015; Norman et al, 2016; Broccard et al, 2017; Mishchenko et al, 2018). Hybrid circuits are typically implemented through a dynamic clamp protocol that injects current computed by a model from an instantaneous voltage recording (Robinson and Kawai, 1993; Sharp et al, 1993; Prinz et al, 2004; Destexhe and Bal, 2009; Nowotny and Varona, 2014). The model can be a simple conductance description or the outcome of a complex biophysical neuron or network simulation. Neuron models used to build hybrid circuits expand from simplified nonlinear equations to Hodgkin-Huxley type paradigms. Synapse models range from standard Ohm’s law implementations for gap junctions to nonlinear graded synapse models that require preand post-synaptic voltage information. Each model has its own set of parameters and they all require specific adaptations for their use in hybrid circuit preparations.

In spite of the large applicability of hybrid circuits to study neuron and network dynamics, including plasticity and learning mechanisms, their use has been somehow limited by the difficulty of their implementation. Hybrid circuit construction often requires specific hardware and/or soft or hard real-time software technology to accurately implement the associated recording and stimulation cycles (Christini et al, 1999; Pinto et al, 2001; Muñiz et al, 2005; Arsiero et al, 2007; Muñiz et al, 2009; Kemenes et al, 2011; Nowotny and Varona, 2012; Linaro et al, 2015; Patel et al, 2017).

Additionally, the construction of hybrid circuits involves handling the different time and amplitude scales of living and model neurons and synapses. In the case of electrophysiological experiments, electrode resistance specifications are also an important issue. All amplitude and time scale adjustments have to be addressed specifically for each preparation. These tasks are time-consuming and often a main source of frustration when building hybrid circuits.

Furthermore, when using hand-tuned parameters in a hybrid configuration, experimentalists frequently have to deal with voltage drift and the natural evolution of membrane potential oscillations during minutes or hours of experimental work. In this paper, we present a set of algorithms to facilitate the building of hybrid circuits using software neurons and synapses. These algorithms perform automatic calibrations and dynamic adaptations of time, voltage amplitude/offset, and current scales to implement open- and closed-loop interactions with living neurons in real time following the scheme illustrated in Fig. 1. Our validation experiments show that the use of these algorithms contributes to better tuned and more natural interactions between living and model neurons, to a reduction of the time expended on adjustments and, thus, to expand the life expectancy of the preparations. We also show that the proposed dynamic adaptation approach for building hybrid circuits is useful to automate experiments, achieve goal-driven control of neural activity, and explore and map neural dynamics. To favor their dissemination and use, the algorithms described in this paper have been implemented in RTHybrid, an open-source library that can be run over different linux platforms, with or without real-time (Amaducci et al, 2017). The proposed algorithms can also be easily migrated to other existing software platforms.

**Figure 1.**
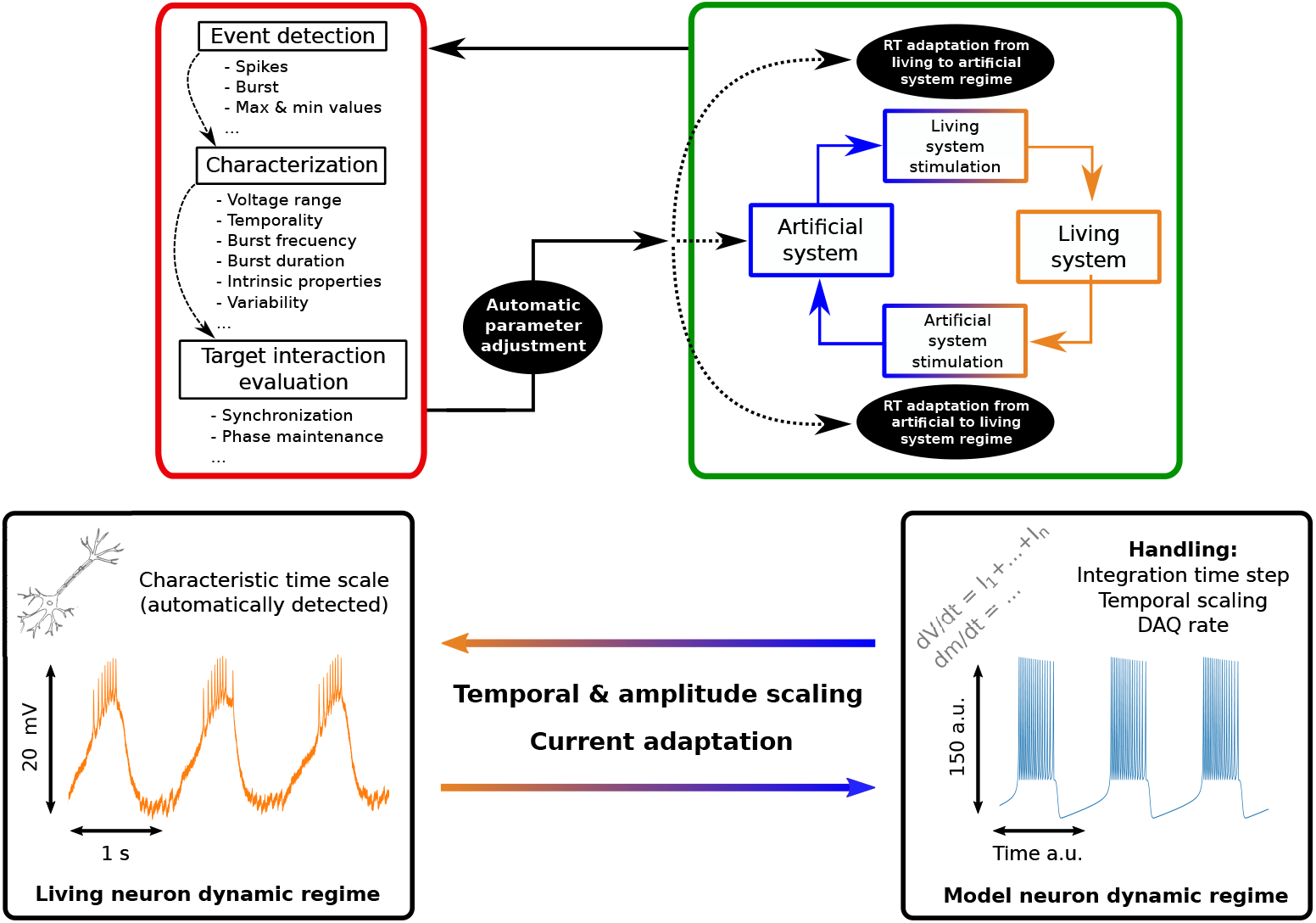
General closed-loop approach proposed for the assisted automatic adaptation in hybrid circuits. Living and model systems are connected mono- or bi-directionally with the required adaptation in each direction to make interaction signals compatible with their dynamical regime in real time. During a test interaction, the system is evaluated through the detection of events that are relevant, e.g. spikes, bursts, maximum and minimum voltage amplitudes, etc. Hybrid circuit efficiency measures for the system characterization are calculated and parameters are adjusted accordingly. Several cycles in the interaction are typically used to assess the hybrid circuit target goal. Adaptation is done in both directions in the case of bidirectional interactions.

## 2 Material and methods

### 2.1 Experimental setup

To validate the automatic adaptation algorithms described in this paper, we built hybrid circuits using the crustacean pyloric central pattern generator (CPG) (Marder and Calabrese, 1996; Selverston, 2005). The neurons of this circuit can be easily identified by their waveform and are particularly resistant to long recordings while sustaining their characteristic spiking/bursting rhythm. As we illustrate in our validation experiments, these neurons generate sequential activity in hybrid circuit experiments using bidirectional interactions with a wide variety of artificial neuron models and synapses ranging from simplified descriptions to conductance-based paradigms.

Adult shore crabs (*Carcinus maenas*) were used in all preparations. They were purchased locally and maintained in a tank with artificial seawater at 13-15°C. Crabs were anesthetized by ice for 15 min before dissection. The procedures followed the European Commission and Universidad Autónoma de Madrid animal treatment guidelines. The stomatogastric nervous system was dissected following standard procedures and pinned in a Sylgard-coated dish containing *Carcinus maenas* saline (in *mM*: 433 *NaCl*, 12 *KCl*, 12 *CaCl*_2_ *·* 2*H*_2_*O*, 20 *MgCl*_2_ *·* 6*H*_2_*O*, 10 *HEP ES*, adjusted to pH 7.60 with 4 M 287 *NaOH*). After desheathing the stomatognathic ganglion (STG), neurons were identified by their membrane potential waveforms and the spikes times observed in extracellular recordings from the corresponding motor nerves. Membrane potential was recorded using 3 M KCl filled microelectrodes (50 *M Ω*) and a DC amplifier (ELC-03M, NPI). Current injection to implement the hybrid connection was delivered with a second electrode on the same neuron. Data was acquired using a A/D board (PCI-6251, National Instruments).

The algorithms were implemented in RTHybrid (www.github.com/GNB-UAM/RTHybrid). For the validation test, we used a computer with an Intel Core i7-6700 processor running Debian 9 under Linux 4.9 with PREEMPT RT patch. Algorithms and RTHybrid were also tested in Ubuntu with Xenomai 3.

### 2.2 Neuron and synapse models

Different neuron model paradigms can be used in hybrid circuit experiments as long as their equations can be integrated in real-time at the desired acquisition rate to implement realistic interactions with living neurons. It is important to note that each model has typically its own time and amplitude scales. The description of many simplified models commonly used considers arbitrary units both for amplitude and time. Other more realistic descriptions, such as conductance-based models, use physiological units. Independently of the time units of the model, the acquisition rate determines the time interval available for the model integration at each interaction cycle. Thus, models need to be implemented taking into account the living neuron time scale, the acquisition frequency and the model integration step.

To validate our approach, in this paper we used different software neuron models with intrinsic rich dynamics and increased complexity, and thus increased computational cost: (i) the Rulkov map (Rulkov, 2002), a two-dimensional iterated map that can display spiking-bursting behavior; (ii) the Izhikevich model, a two-dimensional system of ordinary differential equations with a quadratic voltage nonlinearity and an auxiliary after-spike resetting mechanism (Izhikevich, 2003); (iii) the Hindmarsh-Rose model, a three-dimensional system of ordinary differential equations with cubic nonlinearities (Hindmarsh and Rose, 1984); and (iv) a conductance based model with the characteristic sigmoid voltage dependencies of Hodgkin-Huxley type descriptions (Ghigliazza and Holmes, 2004). In the experiments described below, we employed a chemical graded synapse description frequently used in CPG studies (Golowasch et al, 1999), and also a bidirectional electrical synapse model (Varona et al, 2001). For the integration of the differential equations a (6)5 Runge-Kutta (Hull et al, 1972) numerical method was used. Our results can be generalized to any neuron and synapse models whose equations can be integrated in the hybrid-circuit interaction cycle.

### 2.3 Real-time software technology

Living neurons are not fast computation units, but they can be very precise at the millisecond range. To achieve temporally precise interactions between living and model neurons in a hybrid circuit configuration using a general purpose operative system, real time patches are needed. Following the idea of providing easy to install and implement technology, the proposed algorithms have been developed in RTHybrid (Amaducci et al, 2017) under a real-time operative system, which allows the implementation of closed-looop interactions under millisecond time constraints. The validation experiments shown in this paper were run with a 10 kHz acquisition/stimulation cycle. RTHybrid can be downloaded at www.github.com/GNB-UAM/RTHybrid.

### 2.4 Dynamic clamp hybrid circuit implementation

To implement hybrid circuits in an electrophysiological experiment, it is necessary to record the activity of a presynaptic cell and deliver the corresponding stimulation on a postsynaptic neuron at each step of the acquisition cycle. Typically, this is done using a dynamic clamp protocol to read voltage from the presynaptic neuron and deliver a current into the postsynaptic cell by employing one or multiple electrodes (Destexhe and Bal, 2009). Beyond electrophysiology, a hybrid circuit can use real-time imaging techniques and light (Krook-Magnuson et al, 2013; Prsa et al, 2017) or chemical (Chamorro et al, 2009) stimulation for the implementation of the interaction. The algorithms described in this work aim for automatic adaptation of the signals between living and model networks so that during the interaction they all work in their natural dynamical range. We will illustrate this by implementing the connections with a dynamic clamp protocol in which living neuron voltages are read and adapted to work with synapses that deliver current with the right range both for the living and model neurons at a specified interaction cycle. With the proposed automatic calibration and adaptation protocols, the hybrid circuit is readily implemented in a few seconds regardless of each particular choice for the neuron and synapse models, experimental setup, preparation and/or electrodes.

## 3 Results

### 3.1 Closed-loop approach for automatic calibration and adaptation

The automation of hybrid circuit implementations requires online dynamic signal analysis, precise event detection and real-time model integration. Figure 1 shows our approach to standardize this process. A typical hybrid circuit system consists of mono and/or bidirectional connections between model and living neurons via synaptic models. Table 1 lists the set of algorithms that we have developed for the automatic construction, calibration and evaluation of hybrid circuits. All these algorithms follow the scheme illustrated in the red square of Fig. 1, which is based on real-time event detection, and the dynamic characterization and evaluation of the interaction goal. With this information, online adjustments in the parameters are readily made as a function of predefined performance measurements. At the beginning of the experiment, the living system dynamics are characterized for a few seconds in order to define a first approximation to the needed adaptations. Then, periodically during the experiments, the event detection of the calibration process keeps running to evaluate the interaction in a continuous manner.

**Table 1.**
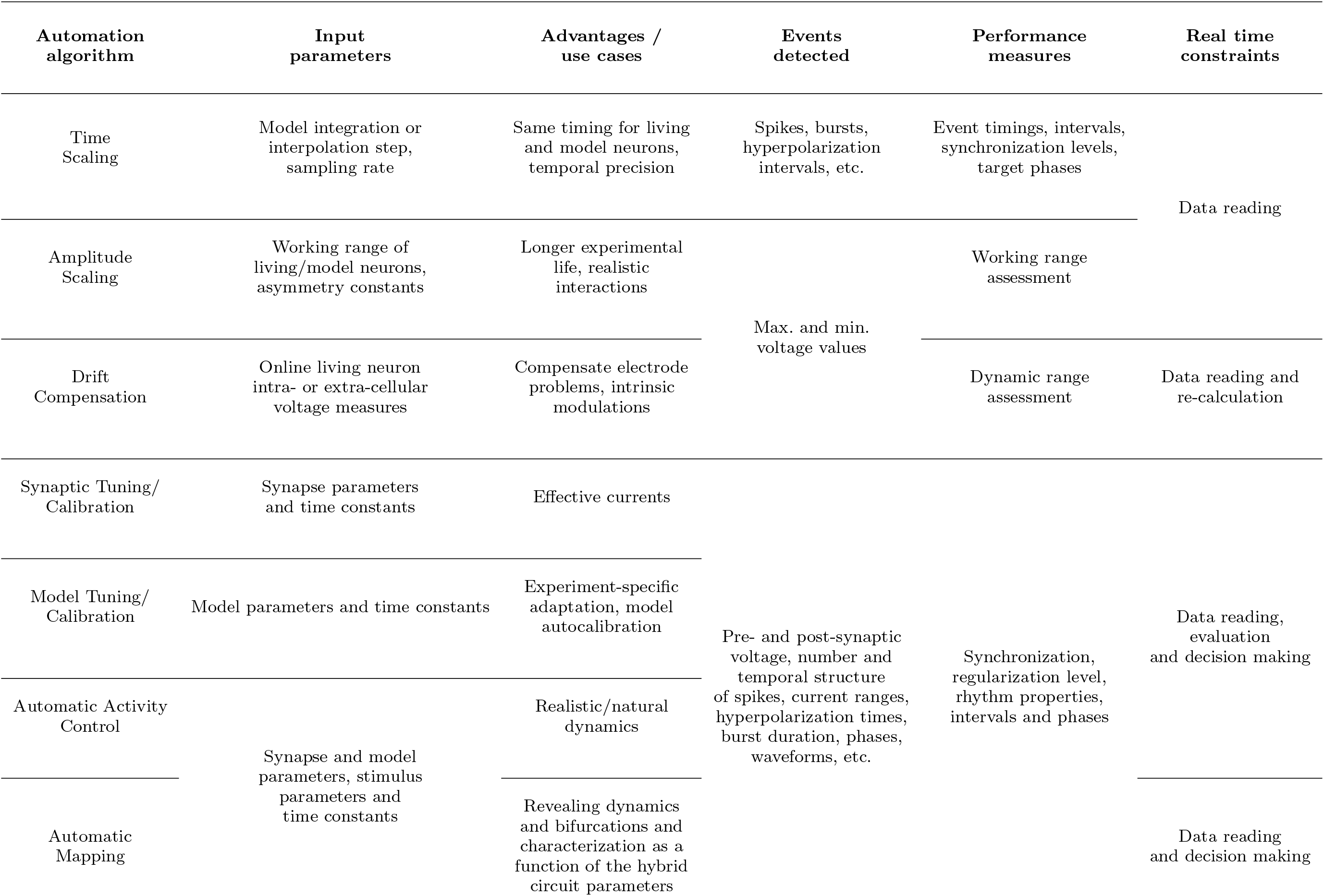
Resume of the different implemented algorithms (rows) and the different requirements, measures, uses cases, etc. (columns) for each one.

### 3.2 Time and amplitude scaling

Since electrophysiological signals and model voltage values have different time and amplitude scales, it is important to note that, at any time, both model and living neurons need to work in their own dynamical range. For this reason, there is a double adaptation so that all neurons receive input in the right range. This allows a natural interaction without causing damage to the living circuit. Thus, the first step in building a hybrid circuit is to conciliate the time and amplitude scales of both living and model networks and the associated synaptic currents. It is important to note that each neuron and synapse model has its own time and amplitude scales. Many models use arbitrary or physiological units with different voltage and time ranges compared to the specific biological counterparts used in a given experiment. The time and amplitude adaptation processes are illustrated in Fig. 2.

**Figure 2.**
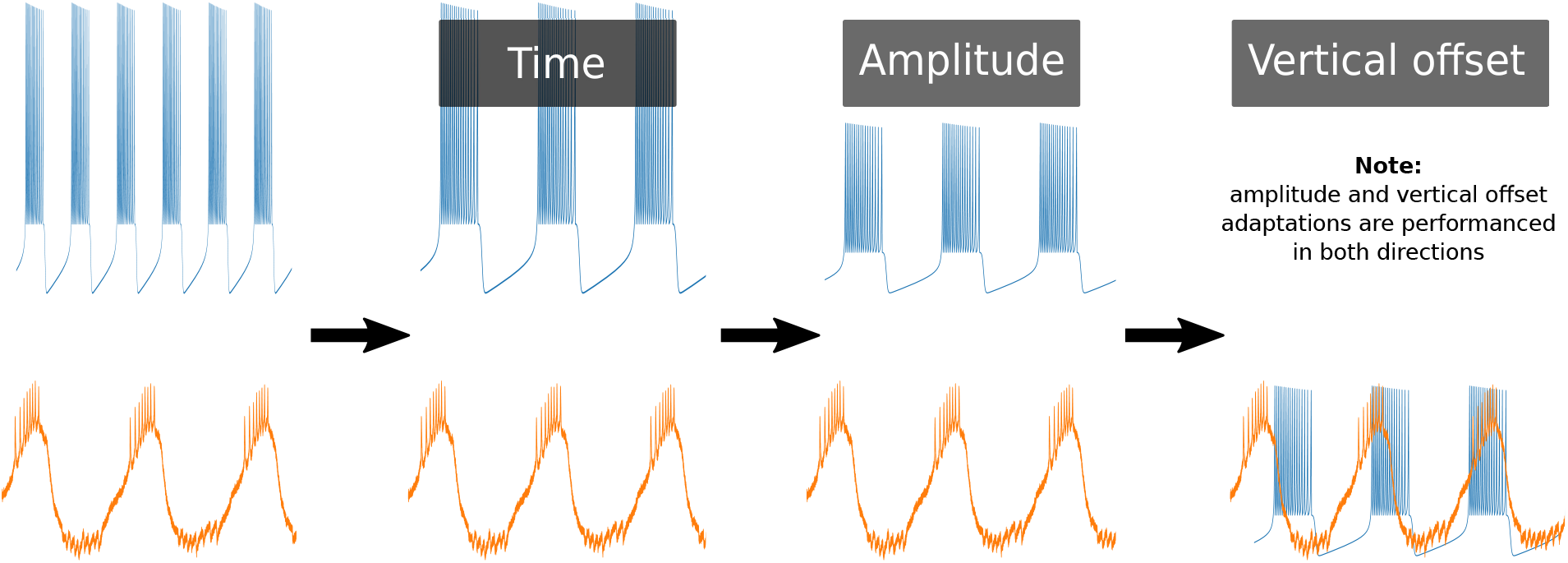
Illustration of the process required to adapt model neurons to a living neuron’s voltage amplitude and time scales, including vertical offset. The right panel corresponds to the resulting adaptation in the working regime of the living neuron.

#### 3.2.1 Time scaling

The algorithm that performs the time calibration of the model takes into account events that define the dynamics of the living neurons under study, such as burst duration, or average period (see Table 1). As illustrated in Fig. 3, different models describe events in their dynamics with distinct time resolution, even when time units are commonly expressed in milliseconds. It is important to note that model integration time (Bettencourt et al, 2008) and the production of values for each dynamic variable in real time are unrelated to the time units of the model. Biophysical models could require many integration steps per acquisition sample to guarantee a precise simulation. Thus, the time scaling algorithm performs a time calibration by measuring the number of points needed to accurately represent reference events in the dynamics of the living neuron, taking into account the model precise integration and the chosen acquisition sampling rate. Reference events that the user can define are, for example, action potentials or bursts (Arroyo et al, 2013; Varona et al, 2016). The sampling rate determines the required discretization of the signals from the living neuron and the model.

**Figure 3.**
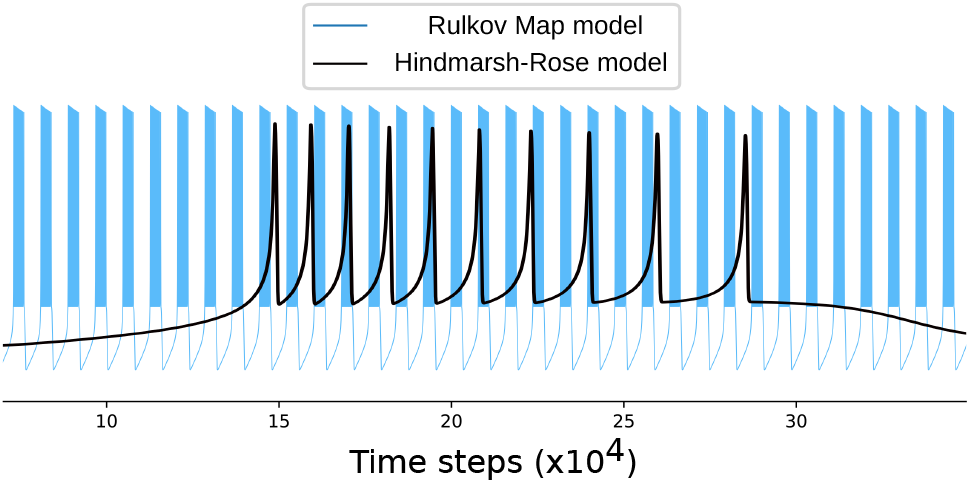
Illustration of different time scales in two neuron models. In this case, the amplitude range is nearly the same but one of the models (the Rulkov map) produces ten bursts while the other (a Hindmarsh-Rose model) produces just one with the same step resolution.

To address this non-trivial task, we adapt the time scale of model neurons and synapses taking into account the characteristic time range of the events defined in the living neuron together with the chosen DAQ sampling rate, which determines the resolution of the discretization. Thus, in our automatic time scaling algorithm, corresponding events (e.g. a burst) are detected both in the signals from the living and model neurons to set the time scale factors. In parallel, information about the DAQ sampling rate and the model minimum integration step are taken into account. With this information two distinct cases can arise:

– If the model has less resolution for an event than the discretization of the living neuron signal set by the sampling rate, the algorithm performs interpolation to provide the required points. This is typically the case of a map model description.
– If the integration of the model generates more points per interaction than needed, the integration step is selected to fulfill two conditions: (i) the time step of the integration has to be smaller than the maximum established for an accurate integration of the model; (ii) the number of points for the reference event in the model must be equal or larger than the sampling rate discretization of the event in the living neurons.

The system also subsamples when the number of points provided by the model integration is larger than those required for the interaction at a particular acquisition rate (see also Table 2). The time scaling algorithm uses real-time detection of predefined reference events from the living neuron signal, such spikes and bursts, to perform the assessments described above.

**Table 2.**
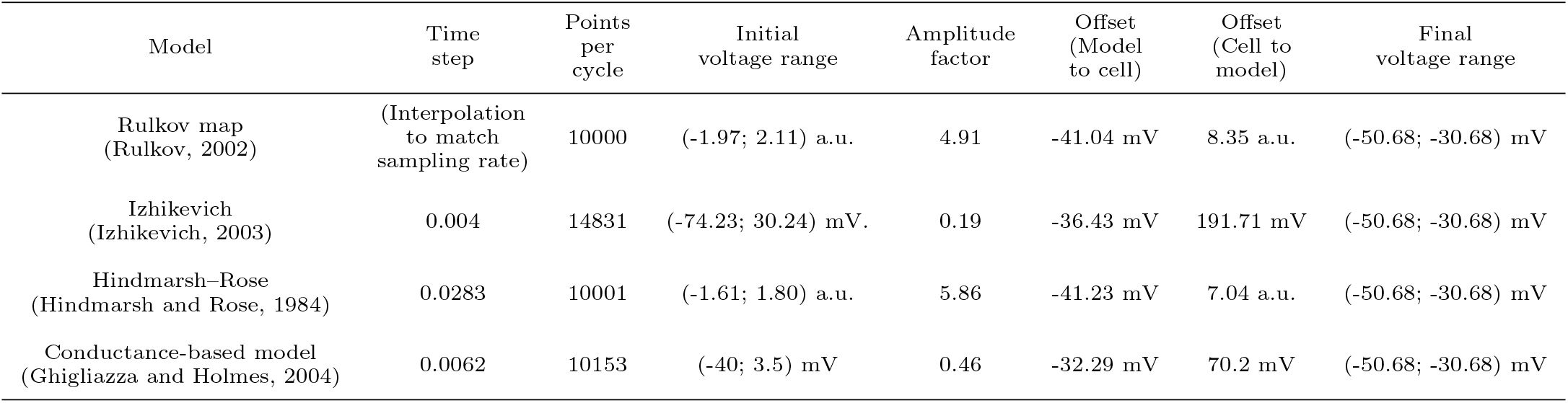
Example of scaling factors for different neuron models to match the time and amplitude of a pyloric neuron that fires approximately one burst per second. Initial values use an online characterization of the living cell dynamics as reference and are re-calculated dynamically during each experiment. The event was a burst and the DAQ frecuency was set to 10 kHz.

#### 3.2.2 Amplitude and offset scaling

Once the time scale has been established, voltage amplitude differences between the living and the model neurons have to be compensated. The parameters of this compensation are typically calculated manually and offline, but this is a slow and tedious process, often taking precious time from the experiment. It is important to note that, even within the same experiment, phenomena like drift or intrinsic changes in the preparation can also produce amplitude and offset changes, which make dynamic voltage compensation necessary.

At any time, each entity of the circuit (living or artificial) must work on its own dynamical range. The adapted voltage value has to be calculated in both directions (model to living and living to model neuron) and used in the synapses to calculate the compensated current to the target neuron. This dynamic adaptation also protects the preparation from excessive current injection, a common issue in manual adaptations that put at risk the living neuron. Figure 4 shows an example of the amplitude adaptation in two neurons that have different voltage scales. These differences consist both on amplitude absolute range and vertical offset.

**Figure 4.**
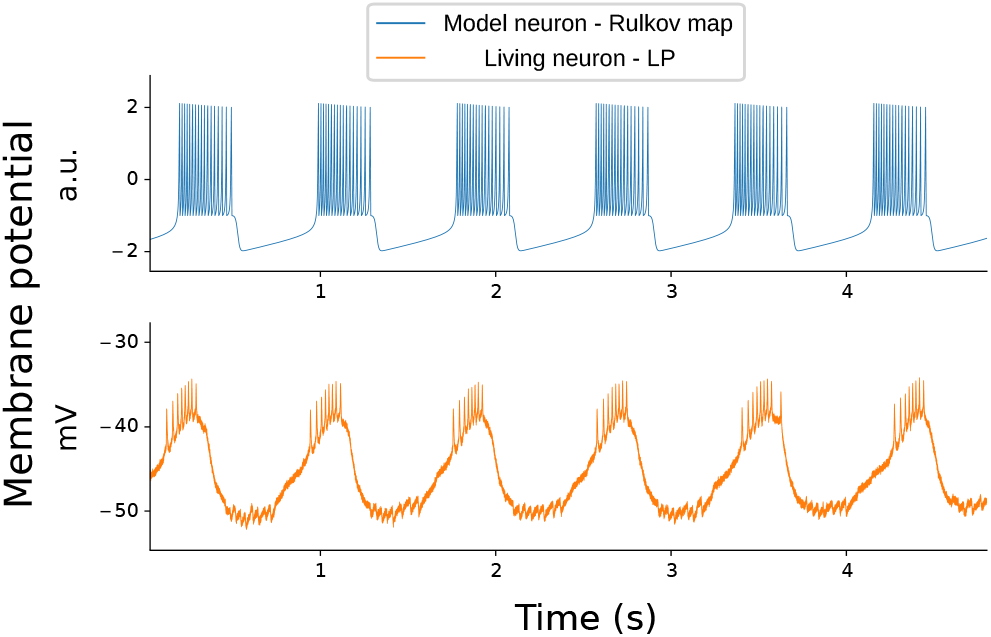
Illustration of the need for voltage amplitude scaling between a model neuron and a living neuron. In this case, the model neuron is a Rulkov map whose bursting amplitude is in the range between -2 and 2 a.u. The living neuron in this example displays bursting activity in the range between -55 to -35 mV. The bursting frequency has already been adapted in the model to match the detected in the living neuron using the time scaling algorithm.

Thus, a second algorithm addresses the characterization of the voltage amplitude range and offset. For this task, a time window is defined in which the algorithm detects the maximun and minimun voltage values of the predefined reference events from the living neuron. With this information, the system calculates a scale amplitude factor and adjusts the vertical offset to match the biological signal range. This is done in both directions, as each neuron continues working on its own dynamical range during the hybrid circuit experiment.

#### 3.2.3 Range-compatible neurons

The joint time and amplitude scaling process is illustrated in Fig. 5. The specific values of the scaling factors automatically obtained for different models when building a hybrid circuit with a pyloric neuron are shown in Table 2.

**Figure 5.**
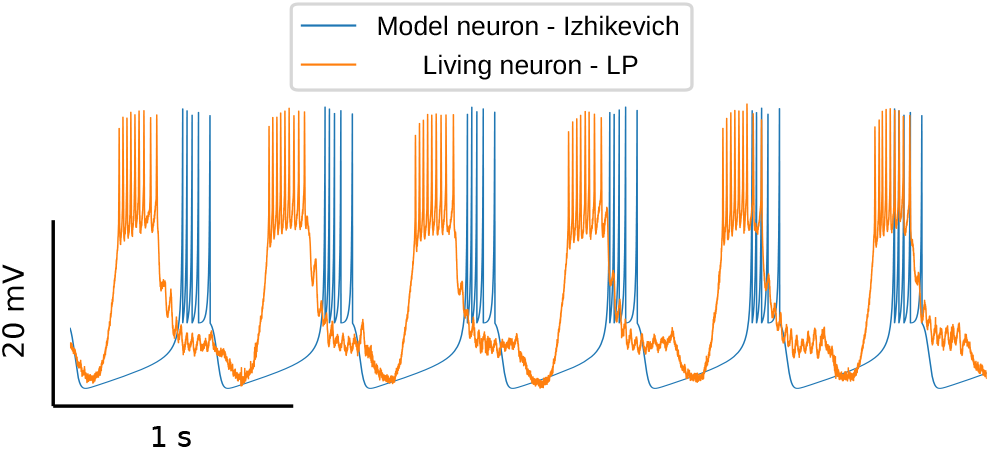
Simultaneous time series of living and model neurons signals after the automatic time and amplitude calibration. This example illustrates the resulting time and amplitude adaptation of a Izhikevich neuron model to work in the dynamical regime of a pyloric LP neuron. At this point there are no synapses between the model and the living neuron, but time and amplitude scale factors are already set and ready to be used in the synapse models.

The automatic processes illustrated in Figs. 3 and 4 occur at the same time and are independently but simultaneously addressed by the calibration algorithms. Figure 5 shows the result of these two adaptations. The time series of both the living cell and the model neuron are plotted together to illustrate that they are in the same time and amplitude scale as in the living neuron control regime. In this example, the synapses of the hybrid circuit are not connected yet but currents in any direction can already use the scaling factors automatically calculated at this point. With both time and amplitude adaptations working at the same time, model neurons are ready to be connected to specific living neurons in a hybrid circuit.

### 3.3 Synapse tuning/calibration

Once amplitude and time scales are compatible between living and model neurons, the connectivity required for the hybrid circuit can be established. Figure 6 shows an example of a hybrid circuit built with a bidirectional connection between the LP neuron of the pyloric CPG and a Izhikevich neuron model (Izhikevich, 2003). The connection models consider one fast and one slow graded sinapses to/from the LP, respectively (Golowasch et al, 1999). Despite the previous adaptations, synapses might also require to adjust parameters such as maximum conductances, time scales and kinetic parameters to achieve the connection goal. For example, a desired behavior for the hybrid circuit can be anti-phase rhythmic activity between the living and the model neuron. This process is not trivial, as synapse parameters may play a key role to establish the target goal. For this goal the synaptic conductances can be increased from a low value until anti-phase bursting behavior is reached.

**Figure 6.**
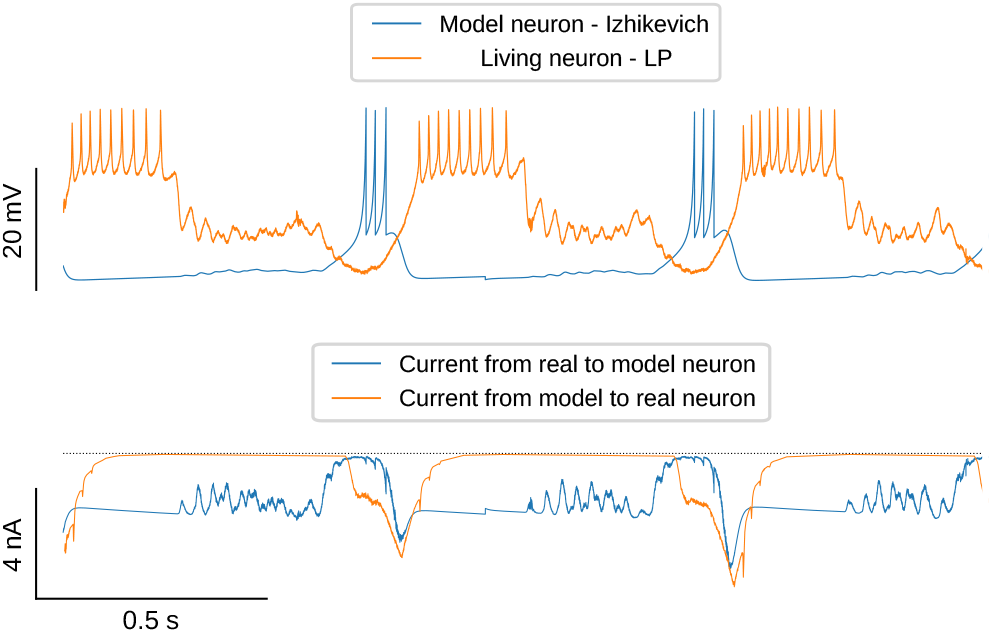
Hybrid circuit between a pyloric LP neuron and a neuron model (Izhikevich, 2003) built with inhibitory graded synapses (Golowasch et al, 1999) that leads to antiphase bursting behavior. Automatic calibration and adaptation of the model and living neuron time and amplitude scales lead to an effective interaction in the hybrid circuit. Top panel shows the living and model neuron voltage signals in the working space of an LP neuron. The bottom panel shows the injected currents. In this example, the synapse to the model neuron is implemented by a fast graded model and the synapse to the living neuron corresponds to a slow graded synapse model.

Thus, to determine the right values of synaptic parameters during the hybrid circuit interaction, a set of goal-driven closed-loop protocols have been designed (for complete needed configurations see Table 1). The general and common structure of these closed-loop protocols consists on three parts (repeated until the connection fulfills the pre-established goal):

i. The first part establishes a continuous event detection during the closed-loop interaction, as described above (see also fig. 1).
ii. Secondly, information is used to characterize and evaluate the performance of the established goal for the interaction with a given metric.
iii. Finally, parameters are changed according to the performance and the goal.

The events for the characterization of the activity, the performance measures for the interaction goal, and the model and synapse parameters must be chosen for of each specific experiment. In this context, characterization measures like frequency, phase, level of activity, etc. can be used for multiple goals, just adjusting the target interaction evaluation. As an example of this process, Fig. 7 shows an experiment where the goal set for the hybrid circuit was the in phase synchronization of a living and a model neuron via a bidirectional electrical synapse. Thus, as a performance measure, the Mean Square Error (MSE) of the two voltage signals was calculated over a time window of three bursts. In this example, the synchronization goal was met when the MSE reached the minimun defined goal. The MSE was measured during five seconds before connecting the two neurons with the electrical synapse. The target MSE was fixed at 40% of the initial MSE value. The conductance of the connection was increased 0.5 mS every three burst cycles until the MSE target value was reached. As can be seen in the figure, this simple protocol leads to the desired synchronization goal. Model parameters can be adjusted analogously, allowing to match the living neuron behaviour, such as the number of spikes in a burst or their temporal structure (Nowotny et al, 2003).

**Figure 7.**
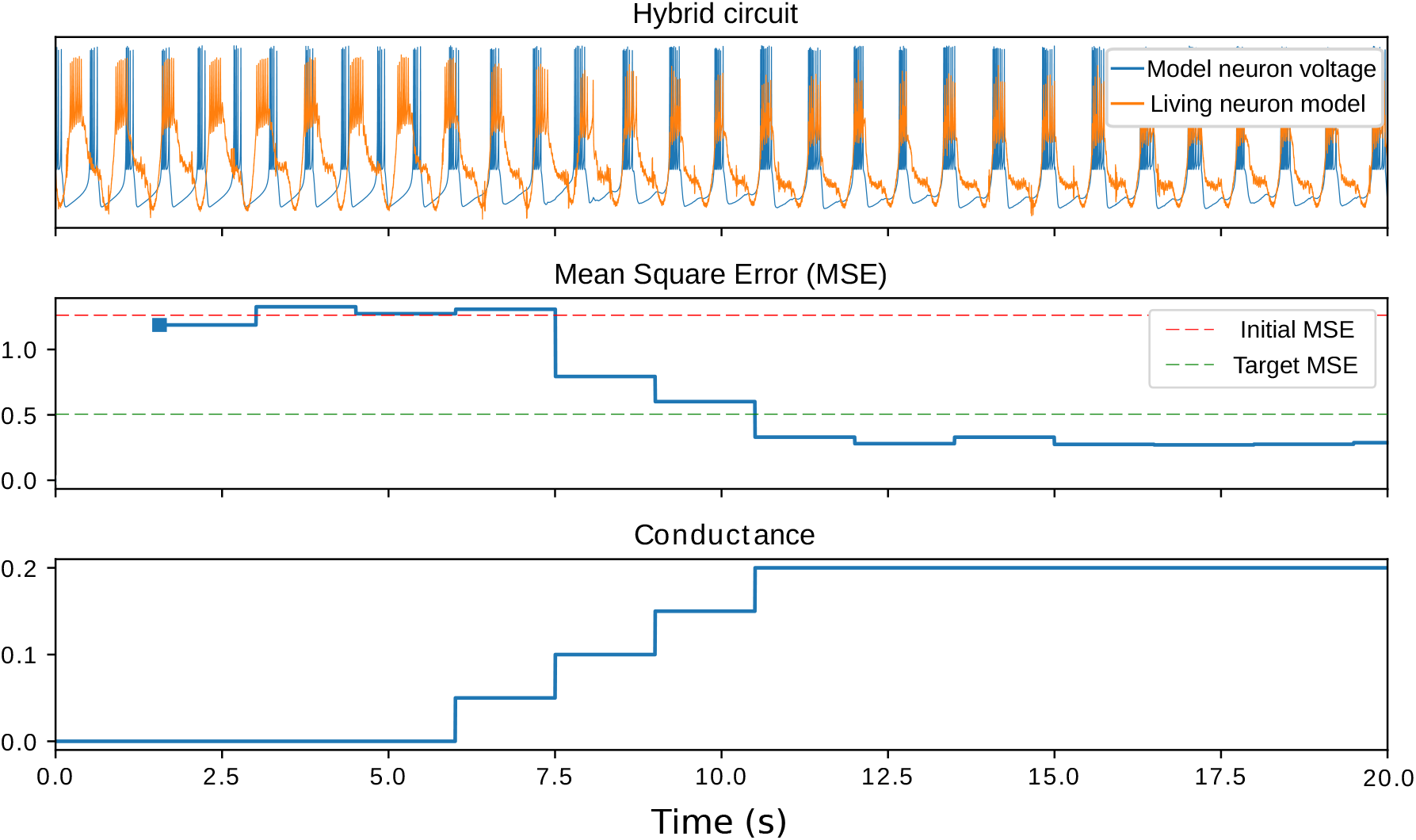
Illustration of the closed-loop synapse calibration process. In this example, a synchronization goal is set to evaluate the efficiency of a hybrid circuit connection. As a performance measurement, the mean square error of the living and model neuron signals is calculated every three bursts. The parameter chosen for the calibration is the conductance of the bidirectional electrical synapse. The top panel shows the evolution of neurons’ membrane potential, middle panel shows the mean square error of the signals and the bottom panel shows the conductance value. This real-time process is also shown in a video included as supplementary material (Online Resource 1).

### 3.4 Drift compensation and ongoing adaptations

Amplitude and time adaptations performed by the previous algorithms might need to be reevaluated periodically during the experiments. This is the case when dealing with voltage drift in the electrodes or a natural evolution of the membrane potential. Continuous monitoring of calibrated ranges is highly relevant, as a change in amplitude or offset can lead to a large current injection into the living neuron or to a malfunction of the hybrid-circuit. Thus, real-time evaluation of voltage ranges and a dynamic adaptation of the drift compensation is also addressed with another algorithm. Figure 8 illustrates this compensation. A time window is defined to set the reevaluation period and the predefined neuron events are again evaluated. Scale factors are thus adjusted as can be seen by the green traces in this figure. In this particular experiment, information was reevaluated every two burst periods and the amplitude factor and vertical offsets were re-calculated accordingly.

**Figure 8.**
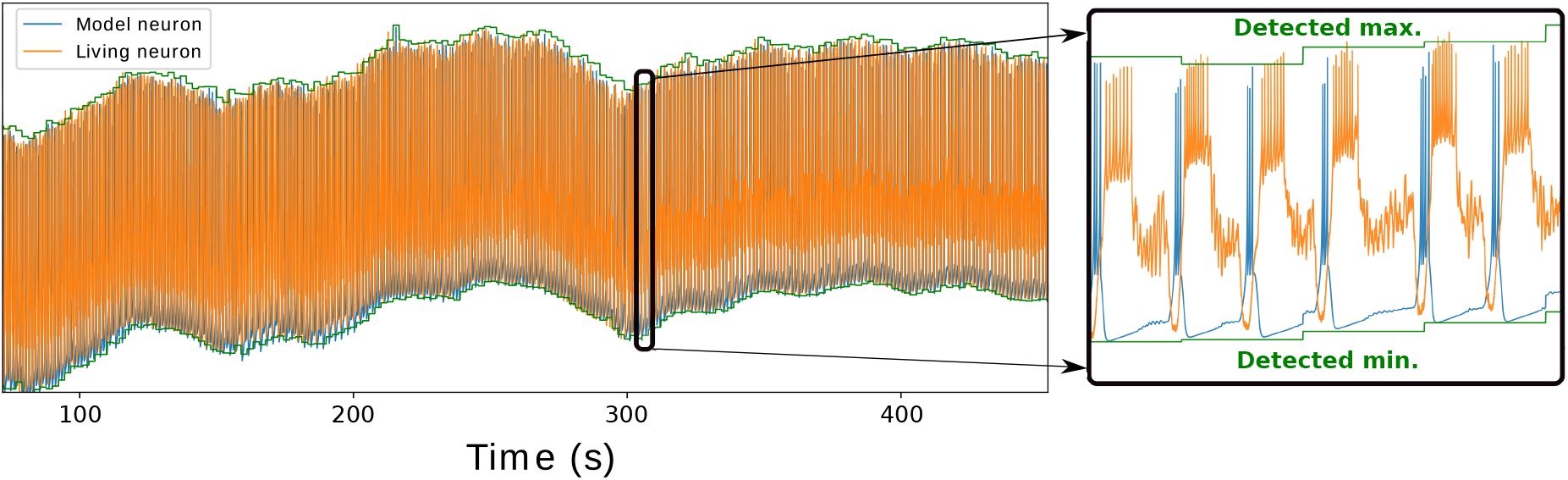
Real-time adaptation to compensate drift during the experiment. Green lines show the maximum and minimum values of voltage amplitude. These values were reevaluated every two cycles of the hybrid circuit interaction based on the activity observed in the living neuron.

### 3.5 Characterization and control of neural dynamics

Hybrid circuits can also be used to characterize and control neural dynamics. For example, interactions with the living system can be used to create new elements in a circuit, to assess the functional role of existing ones, or to replace damaged elements of the living system, while characterizing or sustaining a given dynamics (Szücs et al, 2000; Chamorro et al, 2012; Sakurai and Katz, 2017). A common goal for many hybrid circuits is to reach a certain level of activity, for example a specific regular rhythmic regime (Varona et al, 2001; Hooper et al, 2015).

We illustrate this with a simple goal-driven activity-dependent stimulation algorithm. In this example current is injected into a living LP neuron from the CPG to achieve rhythm regularization. Figure 9 shows how the instantaneous value of the period and its variance is measured within a time window of 5 bursts. This is used as the performance measure for the regularization goal. In control conditions, the living neuron had irregular spiking-bursting activity. The figure shows that once a certain regularization level was reached, further increasing of the current did not lead to a decrease in the variance, and thus to further regularization. Every time the stimulus is turned on or off, the performance measure (in this case the variance) is used to analyze the predefined time window to reevaluate the corresponding change in the signal. In some cases, predefined events for the performance evaluation can be transiently absent and the algorithm waits for their re-appearance. The figure also shows that after the protocol stopped, spiking-bursting irregularity came back. This example illustrates how closed-loop control implemented with an algorithm that tracks events and deals with goal-driven performance measure evaluation allows to achieve the desired result with minimal disturbance to the living circuit.

**Figure 9.**
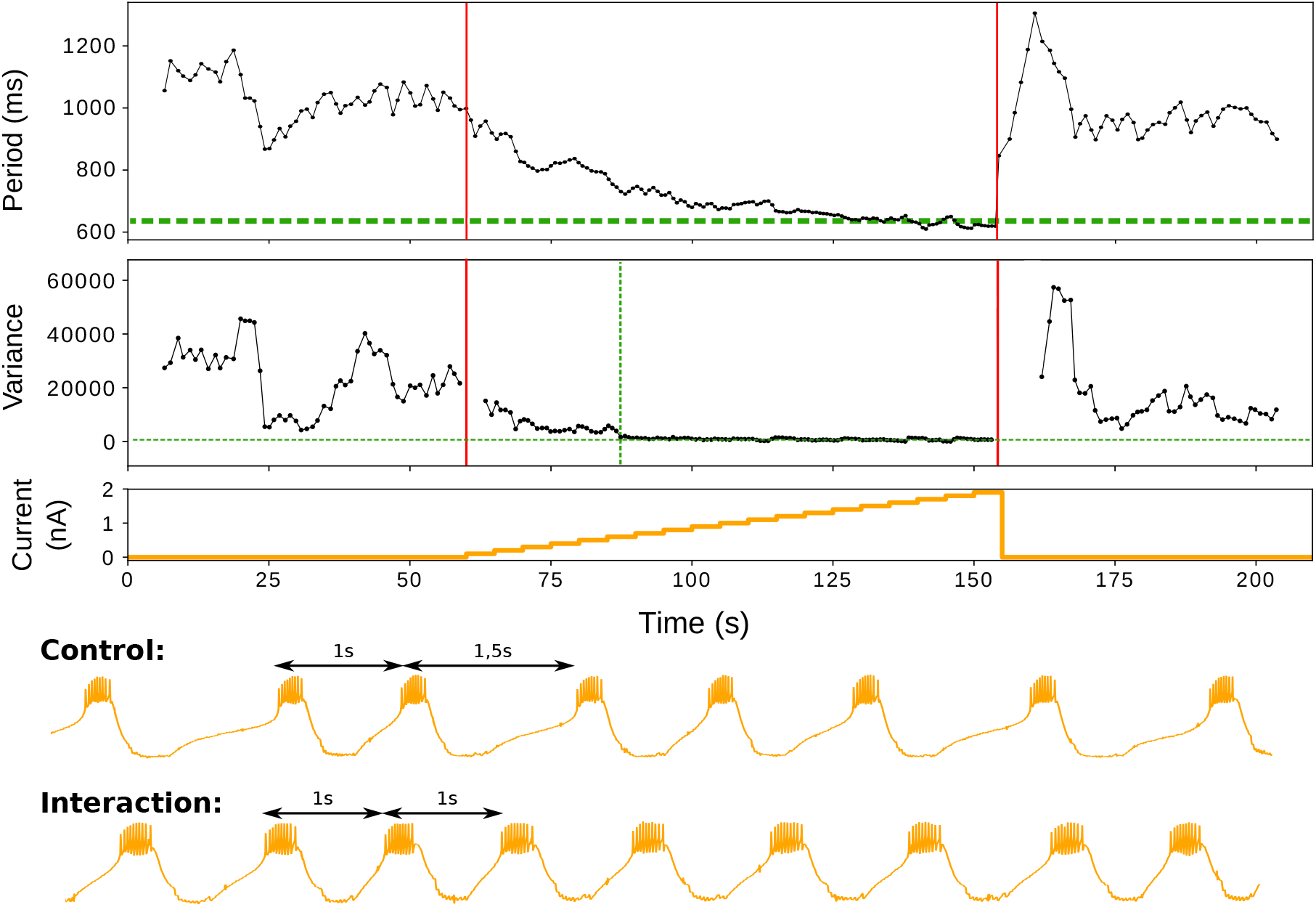
Rhythm regularization of a LP neuron by goal-driven current injection. The recording was monitored in real time, as the system instantaneously measured the period from a predefined event –the burst beginning– and calculated its variance within a time window of 5 bursts. Red lines indicates the start and the end of the current injection. Green dash line in period area indicates minimun value. Horizontal green dash line in variance area indicates zero variance and vertical green line indicates when this value is reached.

### 3.6 Automatic mapping

The same approach discussed in the previous sections can be used to perform automatic parametric searches, and thus to achieve automatic mapping of the hybrid circuit dynamics in relation to a predefined goal. To illustrate the concept of automatic mapping, we developed a protocol to look for a dynamical invariant in a hybrid circuit. A dynamical invariant is defined as a preserved relationship between time intervals that define a sequence in a neural rhythm (Elices et al, 2018). Dynamical invariants are preserved cycle-by-cycle, even during transients.

Thus, we built a hybrid circuit by connecting the LP neuron of the pyloric CPG to a Izhikevich model neuron (Izhikevich, 2003) with a graded chemical synapse (Golowasch et al, 1999). The two parameters to be explored in the mapping of the presence of a dynamical invariant in this circuit were the maximum conductance of the synapse and the threshold for the release of the graded synapse. The presynaptic voltage threshold range was defined as a percentage of the amplitude of the signal. The protocol automatically explored the parameters and built the map for the presence of the invariant.

Figure 10 illustrates the result of this process. For all pairs of parameters shown in the map, the desired antiphase regime was reached, but the dynamical invariant was only present for high correlation (*R*^2^ *>* 90%) values, represented in green. To calculate these values, time events were detected with the tools described in subsection 3.2.1 in order to determine the duration of the intervals defined by the beginning of the burst of the living and the artificial neuron and the instantaneous period of the living neuron. Finally, each *R*^2^ value was calculated with the obtained intervals.

**Figure 10.**
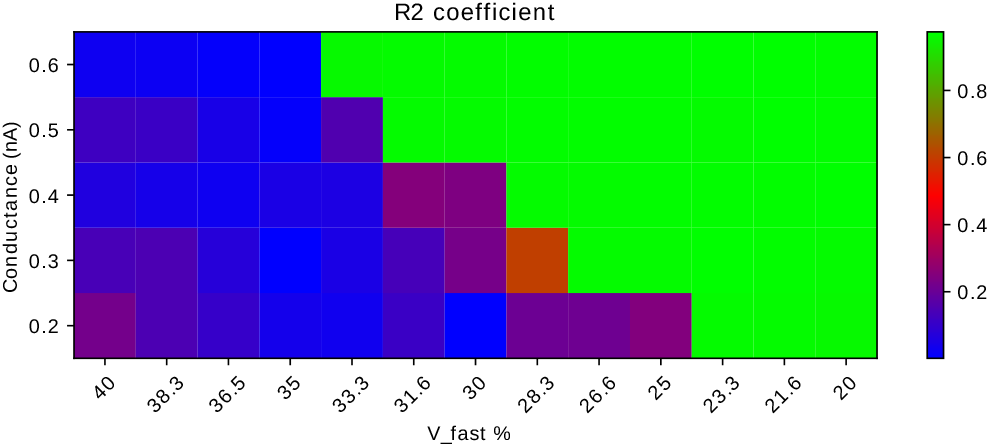
Map of the synaptic parameter space that lead to the presence of a linear relation (a dynamical invariant) between the interval defined by the beginning of the bursting activity of the two neurons (living and artificial) and the instantaneous period of the sequence in the hybrid circuit. The map represents the *R*^2^ of the linear regression. Green regions correspond to the presence of a dynamical invariant between the living and model neuron.

## 4 Discussion

Hybrid circuits are built by connecting living and model neurons. These circuits have a lot of potential in neuroscience research, but require complex experiment-specific adaptations during their construction to work properly. In particular, parameters related to time and amplitude scaling of the models and of synaptic currents involved in the implementation must be evaluated for each preparation and setup. Typically, these adaptations are performed manually by the researcher and, thus, they are time consuming and often sub-optimal. Difficulties associated with the calibration procedures, together with those related to the real-time neuron and synapse model implementation are a major factor preventing the dissemination of hybrid circuit technology. In this work, we have developed a set of algorithms that address these issues and facilitate the automatic building of hybrid circuits.

The proposed algorithms have a wide range of applications to tune and control the behavior of the living circuit in an automated manner, but also to autonomously map the parameter space to achieve a predefined goal, and in general to explore circuit dynamics. It is important to emphasize that in many experiments hard real-time constraints are needed for an artifact-free implementation of hybrid circuits. The algorithms described in this paper have been implemented and validated in RTHybrid, an open-source platform, and can be easily generalized for other closed-loop interactions with the nervous system. Some of the algorithms can also be employed in model simulations alone to evaluate model candidates to be used in hybrid circuits, see also (Elices and Varona, 2015, 2017).

Beyond electrophyology configurations, hybrid circuits can also be implemented using other alternatives such as optical recording and stimulation protocols, e.g. the ones used in optogenetics (Krook-Magnuson et al, 2013) and in neurotransmitter/neuromodulator microinjection protocols (Chamorro et al, 2012). Our algorithms for time and amplitude scaling of the models and, in general, the automation and calibration of goal-driven close-interactions also apply in any context regarding hybrid-circuits. Only the algorithm related to the amplitude scaling in the direction to the living neurons is specific of dynamic-clamp protocols.

Overall, the algorithms described in this paper and the associated standardized strategy to build hybrid-circuits can lead to the dissemination of the use of this technology, contribute to expand the life expectancy of the preparations and favor a new trend in automation of experimental work in neuroscience research.

## Acknowledgements

This work was supported by MINECO/ FEDER DPI2015-65833-P, TIN2014-54580-R, TIN2017-84452-R and ONRG grant N62909-14-1-N279.

## References

Amaducci R, Muñiz C, Reyes-Sanchez M, Rodriguez F, Varona P (2017) On the need for standardized real-time software technology in closed-loop neuroscience. BMC Neuroscience 18:P104

Arroyo D, Chamorro P, Amigó J, Rodríguez F, Varona P (2013) Event detection, multimodality and nonstationarity: Ordinal patterns, a tool to rule them all? The European Physical Journal Special Topics 222(2):457–472, DOI 10.1140/epjst/e2013-01852-9

Arsiero M, Lüscher HR, Giugliano M (2007) Real-time closed-loop electrophysiology: towards new frontiers in in vitro investigations in the neurosciences. Archives italiennes de biologie 145(3):193–209

Bettencourt JC, Lillis KP, Stupin LR, White JA (2008) Effects of imperfect dynamic clamp: Computational and experimental results. Journal of Neuroscience Methods 169(2):282–289, DOI 10.1016/J. JNEUMETH.2007.10.009

Broccard FD, Joshi S, Wang J, Cauwenberghs G (2017) Neuromorphic neural interfaces: from neurophysiological inspiration to biohybrid coupling with nervous systems. Journal of neural engineering 14(4):41,002, DOI 10.1088/1741-2552/aa67a9

Brochini L, Carelli PV, Pinto RD (2011) Single synapse information coding in intraburst spike patterns of central pattern generator motor neurons. Journal of Neuroscience 31(34):12,297–12,306

Chamorro P, Levi R, Rodríguez FB, Pinto RD, Varona P (2009) Real-time activity-dependent drug microinjection. BMC Neuroscience 10(1):P296

Manuel Reyes-Sanchez et al. Chamorro P, Muñiz C, Levi R, Arroyo D, Rodríguez FB, Varona P (2012) Generalization of the dynamic clamp concept in neurophysiology and behavior. PLoS ONE 7(7), DOI 10.1371/journal.pone.0040887

Christini DJ, Stein KM, Markowitz SM, Lerman BB (1999) Practical Real-Time Computing System for Biomedical Experiment Interface. Annals of Biomedical Engineering DOI 10.1114/1.185

Destexhe A, Bal T (2009) Dynamic-Clamp: From Principles to Applications. From Principles to Applications 1:443, DOI 10.1007/978-0-387-89279-5

Elices I, Varona P (2015) Closed-loop control of a minimal central pattern generator network. Neurocomputing 170:55–62, DOI 10.1016/j.neucom.2015.04.097

Elices I, Varona P (2017) Asymmetry Factors Shaping Regular and Irregular Bursting Rhythms in Central Pattern Generators. Frontiers in computational neuroscience 11

Elices I, Levi R, Arroyo D, Rodriguez FdBB, Varona P (2018) Robust dynamical invariants in sequential neural activity. bioRxiv DOI 10.1101/379909

Ghigliazza RM, Holmes P (2004) Minimal Models of Bursting Neurons: How Multiple Currents, Conductances, and Timescales Affect Bifurcation Diagrams. SIAM Journal on Applied Dynamical Systems DOI 10.1137/030602307

Golowasch J, Casey M, Abbott LF, Marder E (1999) Network stability from activity-dependent regulation of neuronal conductances. Neural Computation 11(5):1079–1096, DOI 10.1162/089976699300016359

Grashow R, Brookings T, Marder E (2010) Compensation for variable intrinsic neuronal excitability by circuit-synaptic interactions. Journal of Neuroscience 30(27):9145–9156

Hindmarsh JL, Rose RM (1984) A model of neuronal bursting using three coupled first order differential equations.

Hooper RM, Tikidji-Hamburyan RA, Canavier CC, Prinz AA (2015) Feedback Control of Variability in the Cycle Period of a Central Pattern Generator. Journal of Neurophysiology 114(5):jn.00,365.2015, DOI 10.1152/jn.00365.2015

Hull TE, Enright WH, Fellen BM, Sedgwick AE (1972) Comparing numerical methods for ordinary differential equations. SIAM Journal on Numerical Analysis 9(4):603–637

Izhikevich E (2003) Simple model of spiking neurons. IEEE Transactions on Neural Networks 14(6):1569–1572, DOI 10.1109/TNN.2003.820440, ArXiv

Kemenes I, Marra V, Crossley M, Samu D, Staras K, Kemenes G, Nowotny T (2011) Dynamic clamp with StdpC software. Nature protocols 6(3):405–417

Krook-Magnuson E, Armstrong C, Oijala M, Soltesz I (2013) On-demand optogenetic control of spontaneous seizures in temporal lobe epilepsy. Nature communications 4:1376, DOI 10.1038/ncomms2376

Le Masson G, Renaud-Le Masson S, Debay D, Bal T (2002) Feedback inhibition controls spike transfer in hybrid thalamic circuits. Nature 417(6891):854–858

Linaro D, Couto J, Giugliano M (2015) Real-time Electrophysiology: Using Closed-loop Protocols to Probe Neuronal Dynamics and Beyond. JoVE (Journal of Visualized Experiments) pp e52,320—-e52,320, DOI 10.3791/52320

Marder E, Calabrese RL (1996) Principles of rhythmic motor pattern generation. Physiol Rev 76:687–717

Mishchenko MA, Gerasimova SA, Lebedeva AV, Lepekhina LS, Pisarchik AN, Kazantsev VB (2018) Optoelectronic system for brain neuronal network stimulation. PLOS ONE 13(6):e0198,396, DOI 10.1371/journal.pone.0198396

Muñiz C, Arganda S, Rodriguez FD, de Polavieja GG, Varona P (2005) Realistic stimulation through advanced dynamic-clamp protocols. Mechanisms, Symbols and Models Underlying Cognition, Pt 1, Proceedings 3561:95–105

Muñiz C, Rodríguez FB, Varona P (2009) RTBiomanager: a software platform to expand the applications of real-time technology in neuroscience. BMC Neurosci 10(Suppl 1):P49

Norman SE, Butera RJ, Canavier CC (2016) Stochastic slowly adapting ionic currents may provide a decorrelation mechanism for neural oscillators by causing wander in the intrinsic period. Journal of Neurophysiology DOI 10.1152/jn.00193.2016

Nowotny T, Varona P (2012) Dynamic Clamp, Springer Netherlands, Dordrecht, pp 613–621. DOI 10.1007/978-90-481-9751-4 223

Nowotny T, Varona P (2014) Dynamic Clamp Technique. In: Encyclopedia of Computational Neuroscience, Springer New York, New York, NY, pp 1–4, DOI 10.1007/978-1-4614-7320-6 126-2

Nowotny T, Zhigulin VP, Selverston AI, Abarbanel HDI, Rabinovich MI (2003) Enhancement of synchronization in a hybrid neural circuit by spike-timing dependent plasticity. Journal of Neuroscience 23(30):9776–9785, DOI 23/30/9776[pii]

Olypher A, Cymbalyuk G, Calabrese RL (2006) Hybrid Systems Analysis of the Control of Burst Duration by Low-Voltage-Activated Calcium Current in Leech Heart Interneurons. Journal of Neurophysiology DOI 10.1152/jn.00582.2006

Patel YA, George A, Dorval AD, White JA, Christini DJ, Butera RJ (2017) Hard real-time closed-loop electrophysiology with the Real-Time eXperiment In-Automatic adaptation of model neurons and connections to build hybrid circuits with living networks 13 terface (RTXI). PLoS Computational Biology 13(5), DOI 10.1371/journal.pcbi.1005430

Pinto R, Elson R, Szücs A, Rabinovich M, Selverston A, Abarbanel H (2001) Extended dynamic clamp: controlling up to four neurons using a single desktop computer and interface. Journal of Neuroscience Methods 108(1):39–48, DOI https://doi.org/10.1016/S0165-0270(01)00368-5

Pinto RD, Varona P, Volkovskii AR, Szücs A, Abarbanel HD, Rabinovich MI (2000) Synchronous behavior of two coupled electronic neurons. Physical Review E - Statistical Physics, Plasmas, Fluids, and Related Interdisciplinary Topics 62(2):2644–2656, DOI 10.1103/PhysRevE.62.2644,0001020

Prinz AA, Abbott L, Marder E (2004) The dynamic clamp comes of age. Trends in Neurosciences 27(4):218–224, DOI 10.1016/j.tins.2004.02.004

Prsa M, Galiñanes GL, Huber D (2017) Rapid Integration of Artificial Sensory Feedback during Operant Conditioning of Motor Cortex Neurons. Neuron 93(4):929–939.e6, DOI 10.1016/j.neuron.2017.01.023

Robinson HPC, Kawai N (1993) Injection of digitally synthesized synaptic conductance transients to measure the integrative properties of neurons. Journal of Neuroscience Methods DOI 10.1016/0165-0270(93)90119-C

Rulkov NF (2002) Modeling of spiking-bursting neural behavior using two-dimensional map. Physical Review E - Statistical, Nonlinear, and Soft Matter Physics 65(4)

Sakurai A, Katz PS (2017) Artificial synaptic rewiring demonstrates that distinct neural circuit configurations underlie homologous behaviors. Current Biology 27(12):1721–1734.e3, DOI https://doi.org/10.1016/j.cub.2017.05.016

Selverston AI (2005) A neural infrastructure for rhythmic motor patterns. Cellular and molecular neurobiology 25(2):223–244

Sharp AA, O’Neil MB, Abbott LF, Marder E (1993) The dynamic clamp: artificial conductances in biological neurons. Trends in Neurosciences 16(10):389–394, DOI 10.1016/0166-2236(93)90004-6

Szücs A, Varona P, Volkovskii AR, Abarbanel HDI, Rabinovich MI, Selverston AI (2000) Interacting biological and electronic neurons generate realistic oscillatory rhythms. Neuroreport 11(3):563–569, DOI 10.1097/00001756-200002280-00027

Varona P, Torres JJ, Abarbanel HDI, Rabinovich MI, Elson RC (2001) Dynamics of two electrically coupled chaotic neurons: Experimental observations and model analysis. Biological Cybernetics 84(2):91–101, DOI 10.1007/s004220000198

Varona P, Arroyo D, Rodríguez F, Nowotny T (2016) Chapter 6 - online event detection requirements in closed-loop neuroscience. In: Hady AE (ed) Closed Loop Neuroscience, Academic Press, San Diego, pp 81–91, DOI https://doi.org/10.1016/B978-0-12-802452-2.00006-8

Wang S, Chandrasekaran L, Fernandez FR, White JA, Canavier CC (2012) Short conduction delays cause inhibition rather than excitation to favor synchrony in hybrid neuronal networks of the entorhinal cortex. PLoS computational biology 8(1):e1002,306

Yarom Y (1991) Rhythmogenesis in a hybrid system-interconnecting an olivary neuron to an analog network of coupled oscillators. Neuroscience 44(2):263–275

